# Individual-specific associations between behavioral events and song notes shape budgerigar vocal sequences

**DOI:** 10.1101/2024.02.25.581929

**Authors:** Nakul Wewhare, Anand Krishnan

## Abstract

Complex behavioral sequences such as courtship displays are often multimodal, and coordination between modalities is critically important. However, in learned and variable behavioral sequences such as songs, individual variability may extend to multimodal coordination. However, individual variability in complex multimodal sequences and in coordination between distinct behaviors remains underexplored. Here, we report that budgerigars, which continuously learn and modify their complex warble songs, exhibit associations between behavioral events during courtship and song notes. Some associations are unique to individuals, and others are universal across individuals. Additionally, some individuals exhibit more unique associations than others. We also find that birds warbling without courtship emit all notes with broadly similar odds ratios. The presence of individual variability opens up the possibility that social learning may modify the associations between complex sequences of behaviors, and this results in both individual variability and hierarchical differences in the links between courtship or copulatory behaviors and song.

## INTRODUCTION

Animal vocalizations are often arranged into complex temporal sequences [1–4], and their study sheds light into function and coordination in complex behaviors. Complexity of vocal sequences may span multiple hierarchical levels, from individual to group- or species-specific, serving multiple functions from mate attraction to communicating group identity. Social learning introduces further lability, with sequences changing in species that learn throughout their lifetime [5]. Learning increases variability in sexually selected traits [6], with consequences for evolution and speciation [7].

Acoustic sequences, together with visual movements, are important components of multimodal courtship displays [8–12]. Multimodal displays may enhance information transfer by increasing signal efficacy [9]. However, most studies to date have examined either relatively simple sequences or broad patterns of coordination in multimodal sequences [10,11,13,14]. Whether one sequence (for example, movements) can drive sequence structure in another modality remains poorly understood. Further, open-ended learners such as psittaciform birds continuously modify vocalizations to signal group identity [5,15], and the higher-order associations of these changing sequences with courtship displays and associated behaviors remains unexplored. Individually distinct and variable songs may also lead to individual variation in display behaviors, and therefore in associations between specific display related behaviors and song notes (i.e. choreography), but this has not been systematically investigated.

Here, we examined the association between courtship behaviors and song sequences in an open-ended learner, the budgerigar (*Melopsittacus undulatus*). Budgerigars possess a highly complex warble song, primarily studied in context of courtship, that also serves as a signature of group identity [5,15,16]. Specifically, we examined whether courtship behaviors of males were associated with the emission of certain song notes, thus potentially driving the structure of the warble sequence, and whether these associations were stereotyped across birds or individual-specific.

## MATERIALS AND METHODS

### Study animals

All experimental procedures were performed on the Indian Institute of Science Education and Research (IISER) Bhopal campus, and were approved by the Institutional Animal Ethics Committee (protocol no: IISERB/2022/001) in accordance with the guidelines laid out by the Committee for the Purpose of Control and Supervision of Experiments on Animals (CPCSEA, New Delhi). We studied five male budgerigars (*Melopsittacus undulatus*) (henceforth referred to as birds 1, 2, 3, 4, and 5), who occupied a colony with three females and one bird whose sex could not be conclusively determined by cere color. Birds were given ad libitum water and commercially available bird seed and maintained in a room at 25°C, as per a previously published study [5].

### Recording procedure

We recorded songs and courtship displays from male birds in July 2022. Each male bird was placed in a cage within a soundproof acoustic enclosure of dimensions 75×75×75 cm (Newtech Engineering Systems, Bengaluru, India), with the bird to whom it exhibited the most affiliative behavior and singing. Birds remained in the enclosure for 24 hours to habituate them to the experimental paradigm, and were then video-recorded the next day from 8 AM to 6 PM using GoPro Hero 5/7 cameras (GoPro Inc., San Mateo, USA) at 60FPS/720p (Hero 5) or 100FPS/960p (Hero 7), so as to ensure a wide angle view no matter which camera was being used. We additionally recorded audio using Audiomoth recorders [17] at a sampling rate of 44.1 kHz. Each pair was recorded twice with an interval of one week between the recordings. During the period of our study, we did not detect warble songs or displays from females.

### Analysis

In order to understand the relationship between courtship movements and song notes, we first defined a courtship bout as beginning with either a note or a display step, and ending when neither had occurred for 2 seconds. The latter was based on how long a bird typically took to resume a display after changing perch. We selected ten such courtship bouts from each pair per day, one from each hour between 8 AM and 6 PM, giving us 100 courtship bouts in total from five pairs. Simultaneously, we classified warble elements into different note types using a double-blind verification method and categorizations as in previous works [5,16]. We aligned the audio and video by aligning spectrograms of sound from both sources in Raven Pro 1.8 (Cornell Laboratory of Ornithology, Ithaca, NY, USA) to give us the start of the bout in both channels. To obtain the start and end time of each note in the aligned recorder audio, boxes were plotted manually around each note using the spectrogram view in Raven Pro. For video data, we scored all courtship-related behaviors and their start and end times frame-by-frame using BORIS v8.6.5 [18]. A detailed description of each behavior is given in Table 1 (also see Supplementary Video). In addition, we noted periods of “warble only”, defined as no display element occurring for at least one second, but with warble song notes being emitted. After labeling our audio and video data, we obtained two separate time series of notes and behaviors in a courtship bout. We discretized these into segments of 5 milliseconds, shorter than the shortest note type (7 ms), thus ensuring that multiple events did not occur in a single segment.

**Table 1:**
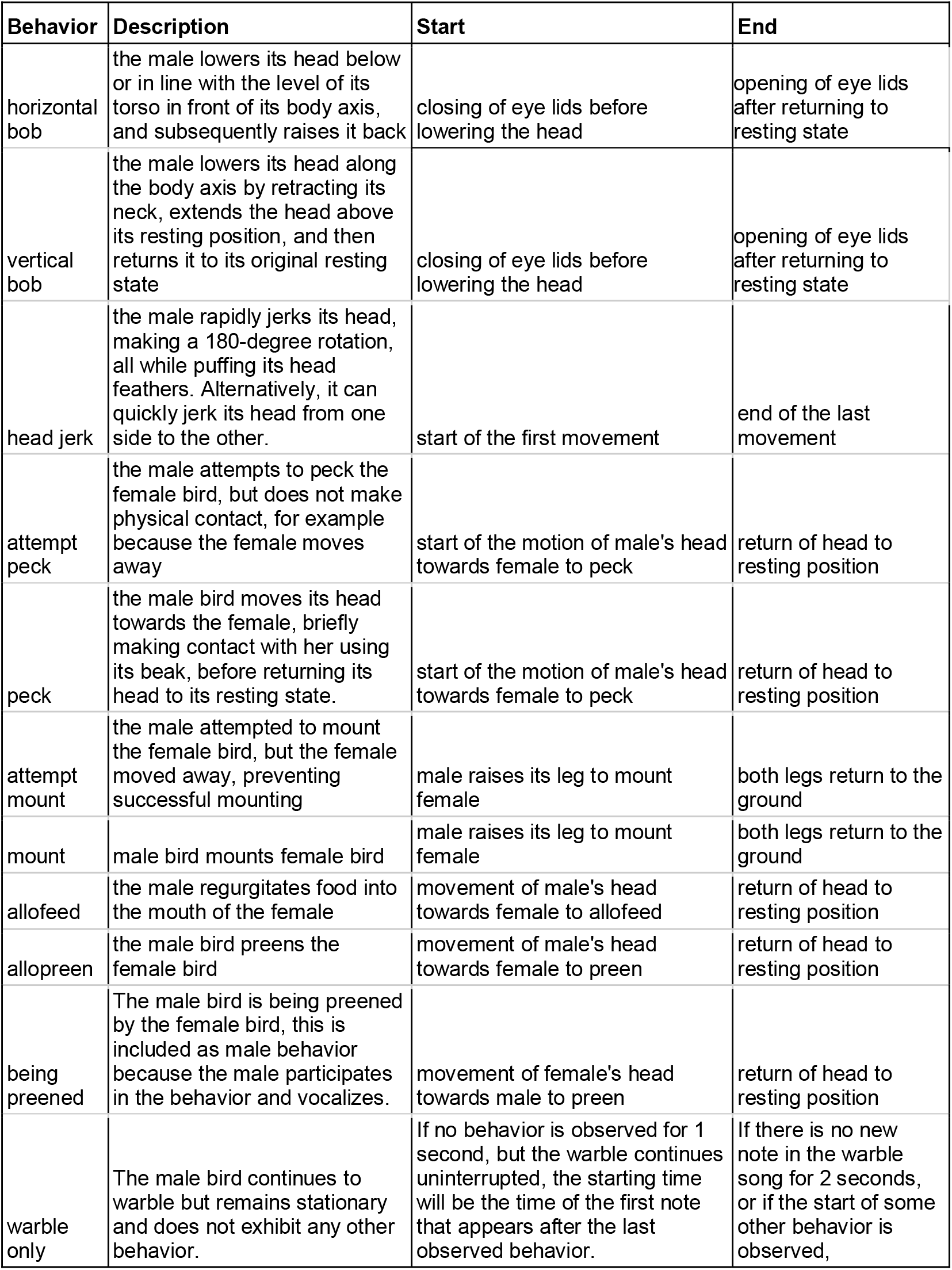
Categories of behaviors analysed from video data, with information on how they were identified and scored. Note: if a display behavior repeats itself again in a series, for example four continuous horizontal bobs, all of them are taken as a horizontal bob event, with start taken as start of first bob and end taken as end of last bob.

Next, we calculated odds ratios of two particular events in the two time-series occurring simultaneously compared to chance. This ratio was the observed number of event co-occurrences divided by the expected number according to chance. The expected value, assuming both time series are independent of each other, was calculated as:

### expected co-occurrences= p(note)^*^p(behavior)^*^l

Where p(note) and p(behavior) represent the probability of a given note and a given behavior in a bout sequence, respectively, and l is the length of the bout sequence.

Next, we calculated the natural log of the odds ratio for all possible combinations of notes and behaviors. Therefore, an odds ratio of 1 resulted in a log-odds of 0, implying that the note and the behavior occurred together as often as expected by chance. A log-odds>0 implied a greater co-occurrence than expected by chance, and vice versa for values <0. To avoid the possibility of rare behaviors exhibiting disproportionately high odds ratios, we calculated the significance of each of our observed log-odds as described below, and only considered significant co-occurrences for further analysis.

First, we randomized our audio and video time series 10,000 times and then calculated log-odds for all combinations of notes and behaviors in each randomized iteration. This gave us a distribution of log-odds for each combination, with known mean and variance based on chance expectations. Next, we calculated the z-score of our observed log-odds based on this randomized distribution, and considered only combinations with significant z-scores for further analysis (all non-significant values were set to zero in visualizations to simplify their interpretation). We considered a “high” odds ratio to be anything higher than one standard deviation above the mean value for all combinations of behavior and notes (considering data from all individual birds combined) (Supplementary Figure 1). The mean log-odds ratio for co-occurrence between song and behavior was -0.08, with a standard deviation of 0.49, resulting in a mean plus standard deviation of 0.41 (i.e. between 0.3-0.45 in a binned histogram) (Supplementary Figure 1). Therefore, to be conservative, we used a value of 0.3 as the threshold to define a “high log-odds ratio” or high co-occurrence, and this threshold encompassed around 1/4th of all possible combinations.

## RESULTS

Our dataset of 100 courtship bouts across 5 pairs consisted of 19,934 annotated song notes and 2,154 courtship-related behavioral events. The bout duration varied from 9-334 seconds (average 54.16 seconds). All five males exhibited high co-occurrence for horizontal bob with the b note, head jerks with a and f notes, and for pecks with the e and f notes (Figure 1) (note categories as detailed in [5]). Additionally, co-occurrence between other behaviors and song notes exhibited patterns specific to certain individual males or groups of males, resulting in a unique pattern of relationships between courtship events and vocal sequences for each bird (Figure 2A, B). Four males (birds 1, 2, 3, and 4) exhibited high co-occurrence of vertical bobs with silent periods (0 in sequences) and head jerks with e notes. Various other combinations of song notes co-occurring with behavioral events during courtship were specific to two or three males, and others to single individuals (Figure 2C). As a result, each individual male had a unique relationship between courtship and the notes comprising the warble song sequence. Two other important patterns stood out here: when warbling without other courtship behaviors (‘warble only’), birds did not exhibit broadly increased associations with any note, largely emitting all notes with similar odds ratios (Figure 1). Second, the number of individual-specific associations varied between birds: some birds (1 and 5) exhibited multiple unique associations between notes and behaviors, whereas others (2 and 4) had very few, and tended to exhibit associations that were also found in other birds (Figure 2C).

**Figure 1:**
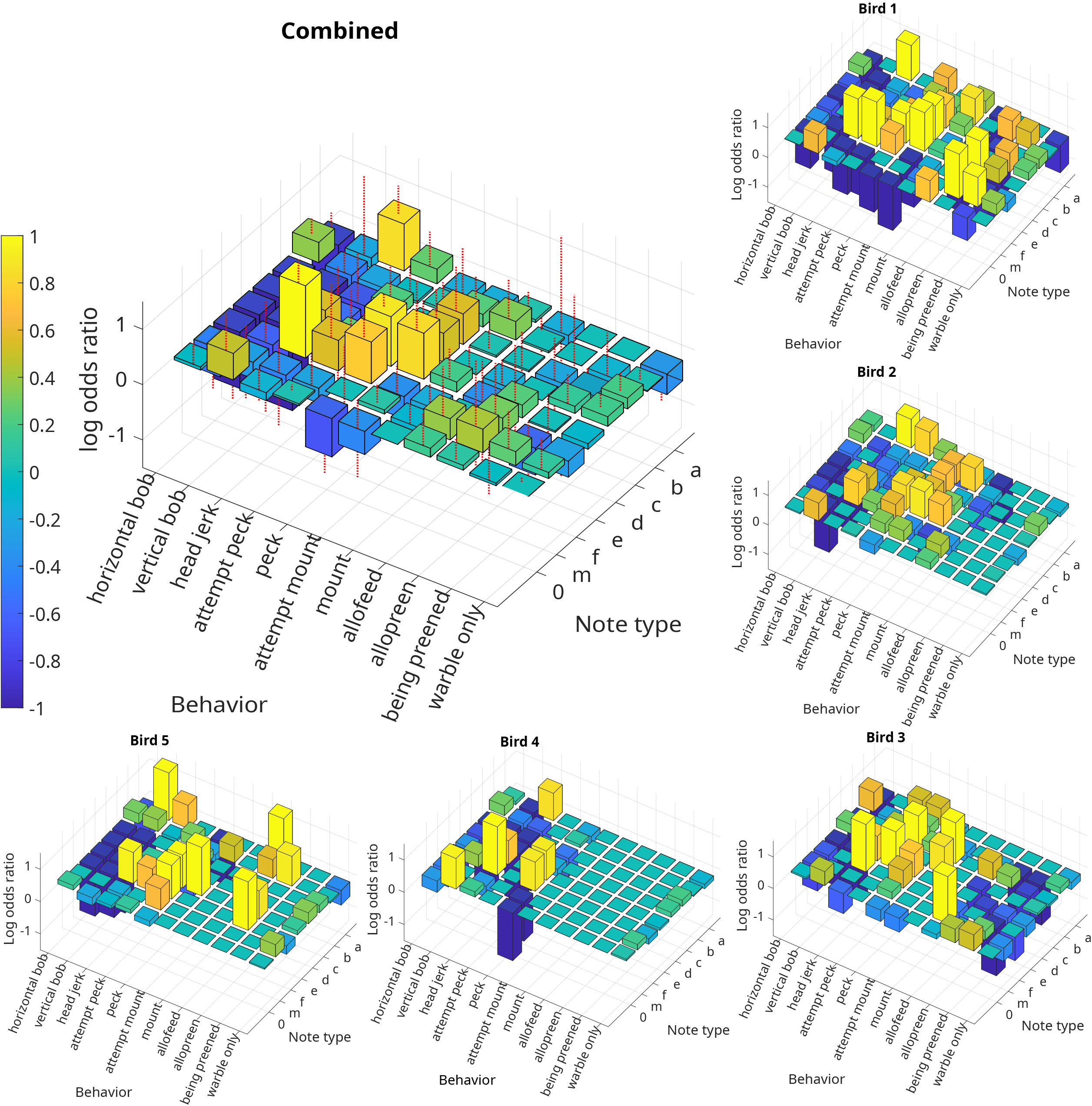
**A**. Three-dimensional histograms showing the strengths of associations (log-odds ratio) between courtship behaviors and song notes for all birds combined (large) and each individual bird (small). Red dashed lines on the large plot indicate standard deviations, and warmer colors indicate higher co-occurrence.

**Figure 2:**
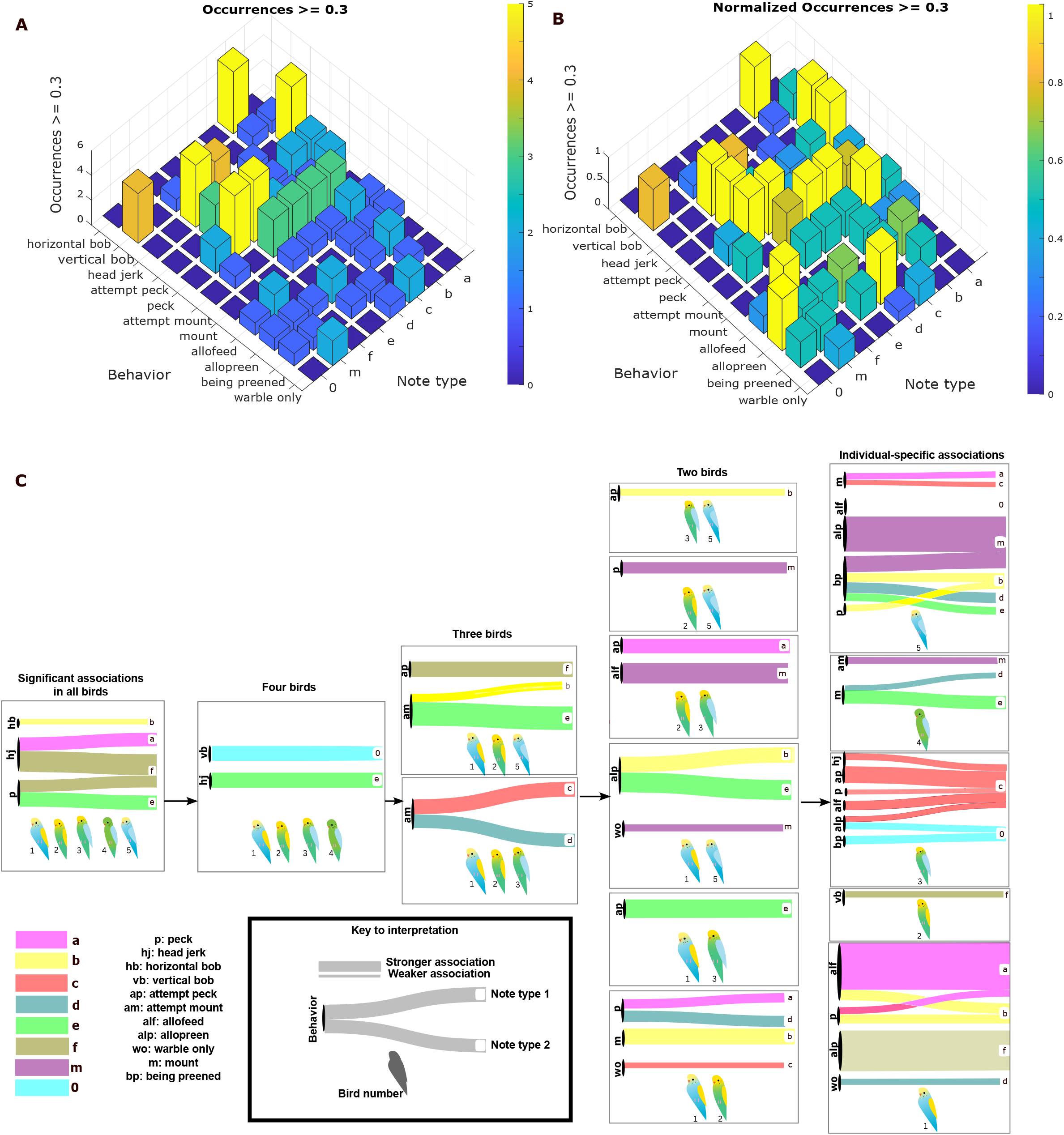
**A**. 3-D histogram showing the number of individuals (vertical axis, 0 to 5) exhibiting a particular song-dance association with high log-odds. **B**. The same plot as in C, normalised to the proportion of individuals exhibiting that association. If three out of five individuals exhibited that particular association, the height of the graph represents the proportion (out of three) where the association was significant (log-odds higher than 0.3). **C**. Each bird shows a unique combination of associations, some found across all birds (left), and others unique to groups of birds or to individuals (right), as indicated by the cartoons at the bottom of each box. The associations are represented as a Sankey diagram, where colors represent the note associated with the behavior (the note is also labelled at the end of each bar, and the behaviors and notes are labelled according to the key in the lower left), and the thickness of the bar is scaled in proportion to the average log-odds value for individuals in which the strength of that particular association crossed the threshold of 0.3. Thus, the thickness of the bar indicates the strength of the association (high log-odds ratio or high co-occurrence). A key to interpreting this figure is also provided. Each cartoon indicates a bird or a group of birds with their numbers, the left side of each box has the abbreviations for each behavior, and the colored boxes correspond to the note that behavior is associated with. Some behaviors may be associated with two or more notes, and thus have two or more bars emerging from them.

## DISCUSSION

Although the relationship between song and courtship display has received study in diverse birds [8–10], our study demonstrates individual variability in these associations. Some individuals have more unique and varied associations between song and behavior than others, potentially indicating the presence of hierarchies that govern courtship and display behavior in social, open-ended learners [5,15]. Although the word choreography is often used to include steps that involve foot or head movements (for example, horizontal or vertical bobs in our data), we also found associations between song notes and other affiliative and copulatory behaviors associated with courtship, such as attempted mounting and allopreening. Further, our study suggests that visual display movements and related behaviors may modify the structure of vocal sequences in a context-dependent manner. In support of this, we note that birds warbling outside of the courtship display (‘warble only’) did not exhibit strong associations with any particular note type. Because budgerigar songs are continually modified by social learning [5,15], it is possible that courtship behavior and associations with song notes are also learned, and may be modified over the course of a bird’s lifetime in response to social factors. This remains to be tested, however, and represents an interesting future line of research.

Warble song serves functions other than courtship, including communicating group identity through higher-order patterns of note repetition embedded in lengthy sequences [5]. The complexity of the budgerigar warble thus serves to highlight its multifunctionality. We propose that a combination of social learning and associations between song and behavior may result in signatures of both individual and colony identity, and that the complexity of these signatures may depend on social structure within a group. Future studies will seek to disentangle these effects and the effect of learning on courtship display [19]. More broadly, however, we highlight how the multifunctional budgerigar warble is a tractable system to study individual variability in complex behaviors, and its relationship to social context.

## ACKNOWLEDGMENTS AND FUNDING

We thank Priya Binwal, Kezia Kunapareddy and Suneel for their help, JNCASR, IISER Bhopal and the Science and Engineering Board (SERB), Government of India (CRG/2022/000187) for funding.

## DECLARATION OF INTERESTS

The authors declare no competing interests.

## FIGURE LEGENDS

**Supplementary Figure 1: A**. Distribution of all log-odds ratios for all associations between song and dance. **B**. Ranked distribution of log-odds, with a cut-off of 0.3 indicating a significantly high log-odds, identified using the mean and standard deviation of all values obtained (see Methods).

## Notes

### Competing Interest Statement

The authors have declared no competing interest.

## REFERENCES

1. Kershenbaum A et al. 2016 Acoustic sequences in non-human animals: A tutorial review and prospectus. Biol. Rev. 91, 13–52. (doi:10.1111/brv.12160)

2. Kershenbaum A, Bowles AE, Freeberg TM, Jin DZ, Lameira AR, Bohn K. 2014 Animal vocal sequences: Not the Markov chains we thought they were. Proc. R. Soc. B Biol. Sci. 281. (doi:10.1098/rspb.2014.1370)

3. Engesser S, Townsend SW. 2019 Combinatoriality in the vocal systems of nonhuman animals. Wiley Interdiscip. Rev. Cogn. Sci. 10, 1–10. (doi:10.1002/wcs.1493)

4. Bhat AS, Sane VA, Seshadri KS, Krishnan A. 2022 Behavioural context shapes vocal sequences in two anuran species with different repertoire sizes. Anim. Behav. 184, 111–129. (doi:10.1016/j.anbehav.2021.12.004)

5. Madabhushi AJ, Wewhare N, Binwal P, Agarwal V, Krishnan A. 2023 Higher-order dialectic variation and syntactic convergence in the complex warble song of budgerigars. J. Exp. Biol. 226. (doi:10.1242/jeb.245678)

6. Lachlan RF, Servedio MR. 2004 Song learning accelerates allopatric speciation. Evolution (N. Y). 58, 2049–2063.

7. Madabhushi AJ, Bhat AS, Krishnan A. 2023 Allopatric montane wren-babblers exhibit similar song notes but divergent vocal sequences. Behav. Ecol. Sociobiol. 77, 1–13. (doi:10.1007/s00265-023-03385-9)

8. Cooper BG, Goller F. 2004 Multimodal Signals: Enhancement and Constraint of Song Motor Patterns by Visual Display. Science (80-. ). 303, 544–546. (doi:10.1126/science.1091099)

9. Ullrich R, Norton P, Scharff C. 2016 Waltzing Taeniopygia: Integration of courtship song and dance in the domesticated Australian zebra finch. Anim. Behav. 112, 285–300. (doi:10.1016/j.anbehav.2015.11.012)

10. Dalziell AH, Peters RA, Cockburn A, Dorland AD, Maisey AC, Magrath RD. 2013 Dance choreography is coordinated with song repertoire in a complex avian display. Curr. Biol. 23, 1132–1135. (doi:10.1016/j.cub.2013.05.018)

11. Miles MC, Fuxjager MJ. 2018 Animal choreography of song and dance: a case study in the Montezuma oropendola, Psarocolius montezuma. Anim. Behav. 140, 99–107. (doi:10.1016/j.anbehav.2018.04.006)

12. Trainer JM, McDonald DB, Learn WA. 2002 The development of coordinated singing in cooperatively displaying long-tailed manakins. Behav. Ecol. 13, 65–69. (doi:10.1093/beheco/13.1.65)

13. Ota N. 2020 Tap dancers in the wild: field observations of multimodal courtship displays in socially monogamous songbirds. Sci. Nat. 107. (doi:10.1007/s00114-020-01686-x)

14. Ota N, Gahr M, Soma M. 2015 Tap dancing birds: The multimodal mutual courtship display of males and females in a socially monogamous songbird. Sci. Rep. 5, 6–11. (doi:10.1038/srep16614)

15. Farabaugh SM, Linzenbold A, Dooling RJ. 1994 Vocal plasticity in budgerigars (Melopsittacus undulatus): evidence for social factors in the learning of contact calls. J. Comp. Psychol. 108, 81–92. (doi:10.1037/0735-7036.108.1.81)

16. Tu HW, Smith EW, Dooling RJ. 2011 Acoustic and Perceptual Categories of Vocal Elements in the Warble Song of Budgerigars (Melopsittacus undulatus). J. Comp. Psychol. 125, 420–430. (doi:10.1037/a0024396)

17. Hill AP, Prince P, Snaddon JL, Doncaster CP, Rogers A. 2019 AudioMoth: A low-cost acoustic device for monitoring biodiversity and the environment. HardwareX 6, e00073. (doi:10.1016/j.ohx.2019.e00073)

18. Friard O, Gamba M. 2016 BORIS: a free, versatile open-source event-logging software for video/audio coding and live observations. Methods Ecol. Evol. 7, 1325–1330. (doi:10.1111/2041-210X.12584)

19. Soma M, Iwama M, Nakajima R, Endo R. 2019 Early-life lessons of the courtship dance in a dance-duetting songbird, the Java sparrow. R. Soc. Open Sci. 6. (doi:10.1098/rsos.190563)

