## Supplementary figures and images for "Individual-specific associations between behavioral events and song notes shape budgerigar vocal sequences"

### Supplementary Figure 1

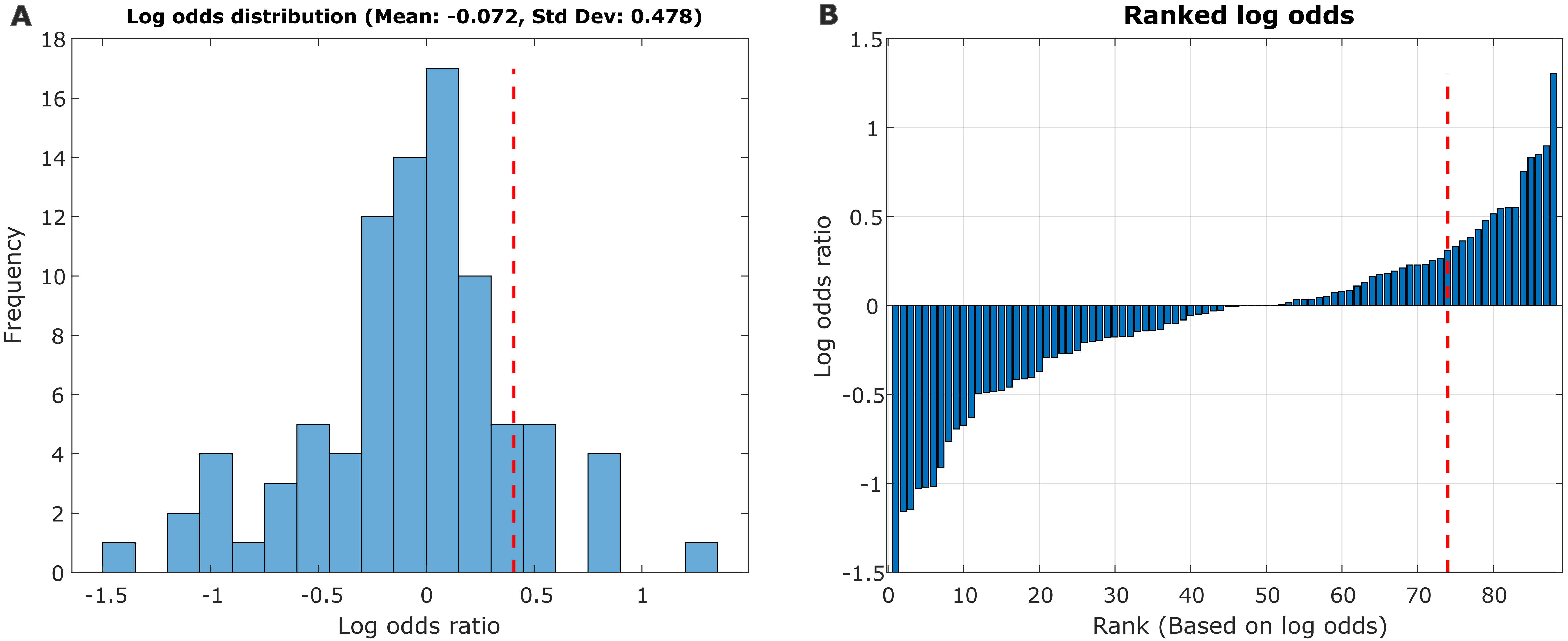
